# Ultrasensitive Proteomics Depicted an In-depth Landscape for Mouse Embryo

**DOI:** 10.1101/2023.01.06.523005

**Authors:** Lei Gu, Xumiao Li, Wencheng Zhu, Yi Shen, Qinqin Wang, Huiping Zhang, Jingquan Li, Ziyi Li, Zhen Liu, Chen Li, Hui Wang

**Affiliations:** Center for Single-Cell Omics, School of Public Health, Shanghai Jiao Tong University School of Medicine, Shanghai, P. R. China, 200025; Institute of Neuroscience, CAS Center for Excellence in Brain Science and Intelligence Technology, CAS Key Laboratory of Primate Neurobiology, State Key Laboratory of Neuroscience, Chinese Academy of Sciences, Shanghai, P. R. China, 200031; Shanghai Applied Protein Technology Co., Ltd., Shanghai, P. R. China, 201100

## Abstract

In recent years, single-cell or low-input multi-omics techniques have brought a revolution in the study of pre-implantation embryo development. However, single-cell or low-input proteome research in this field is relatively underdeveloped, due to the limited source of mammalian embryo samples, the objective reality of high abundance zona pellucida proteins, and the lack of hypersensitive proteome technology. Here, a comprehensive solution of ultrasensitive proteome technology was developed for single-cell or low-input mouse embryos. Both deep coverage route and high-throughput route could significantly reduce the starting material and enhance the proteomic depth without any customized instrument. Using the deep coverage route, an average of 2,665 or 4,585 protein groups can be identified from 1 or 20 mouse zygotes respectively. Using the high-throughput route, 300 single mouse zygotes can be analysis in 8 days with an average of 2,371 proteins identified. With its popularization, we believe researchers can choose deep coverage or high-throughput technology routes according to their own conditions.

## Introduction

Studies on oocyte meiotic maturation and pre-implantation embryo development can provide informative and unique biological insights for mammalian reproduction and development. The process of oocyte meiotic maturation involves the resumption of meiosis and completion of the first meiosis in primary oocytes that arrested in prophase of the meiosis I or germinal vesicle (GV) stage, followed by the initiation of the second meiotic division and arrest in metaphase II (MII) [1,2]. Following fertilization, the embryo undergoes a series of cleavage divisions during pre-implantation development, which encompasses the zygote (or 1-cell), 2-cell, 4-cell, 8-cell, morula, and blastocyst stages [3]. This developmental process is a complex biological stage involving some critical events, such as zygotic genome activation (ZGA), and the first cell lineage differentiation [3,4]. Several recent studies based on single-cell or low-input multi-omics technologies have deciphered this complex and mysterious developmental process at the genetic [5–8], epigenetic [9,10], translational [4,5,11], and metabolic [9,12] levels in hope of obtaining more valuable and full-scale biological insights. Nevertheless, proteins are the actual executors of most biological processes, and gene expression during development requires confirmation at the protein level [3]. There is a growing body of literature that recognizes the poor correlation between transcriptomic and proteomic data at the same developmental stage [13–16]. Taken together, it is necessary to delineate the proteomic landscape of this developmental process.

Mass spectrometry (MS) has become a powerful tool for comprehensive and unbiased proteomic analysis [17,18]. Several studies have explored the proteome of oocyte meiotic maturation and pre-implantation embryo development [13,15,19–25]. Nevertheless, all above studies required thousands of oocytes or embryos as the starting materials to meet the depth of the proteome, which caused hundreds of thousands of animals sacrificing. Therefore, the large number of embryos consumed has discouraged many researchers and limited the development of related fields. The good news is that, with the progress of mass spectrometry and the innovation of single-cell analysis strategies in recent years, it has become possible to illustrate the proteomic profiling of trace or even single cells based on MS. From one mammalian somatic cell, more than 2,000 proteins can be quantified reliably [26–28]. There are also several studies that have applied single-cell proteomic techniques to the oocyte maturation. Guo et al. have achieved the quantification of ∼1,900 proteins on average from one human oocyte [16]. But the size of oocytes differs between mouse and human resulted in a difference in protein content about 4-fold [29], which causes more challenging to illustrate single mouse oocyte proteomic landscapes. Using the ingenious single-cell proteomic strategy nanoliter-scale oil-air-droplet (OAD) chip, Li et al. identified 355 proteins from single mouse oocyte [30]. With the same preparation methods as human oocytes mentioned above, Guo et al. identified no more than 500 protein groups in single mouse oocytes [16]. These proteomic depths have been far from the start line for solving biological problems. It further highlights the scarcity and importance of biomaterials derived *in vivo* and the need to develop new methods to completely characterize the oocyte or embryo proteome.

In the present study, we reported a comprehensive scheme for protein profiling of mouse oocytes and embryos from single-cell to low-input (Fig. 1). Both deep coverage route and high-throughput route could significantly reduce the starting material, and be selected by investigators based on their own demands. In brief, our ultrasensitive proteome platform quantified up to an average of 2,665 protein groups in single zygotes. And when the starting material increased to 20 zygotes, the number of quantified proteins elevated to 4,585 on average. As far as we know, the very early stage of mouse maternal-to-zygotic transition lacks a large-scale proteome landscape using low-input approach. We also established a high-throughput proteome platform based on TMT labelling, which can complete the analysis of 300 single zygotes in ∼8 days with an average of 2,371 proteins identified. This platform raises the possibility of large-scale proteomic researches about oocyte meiotic maturation and pre-implantation embryo development.

**Figure 1.**
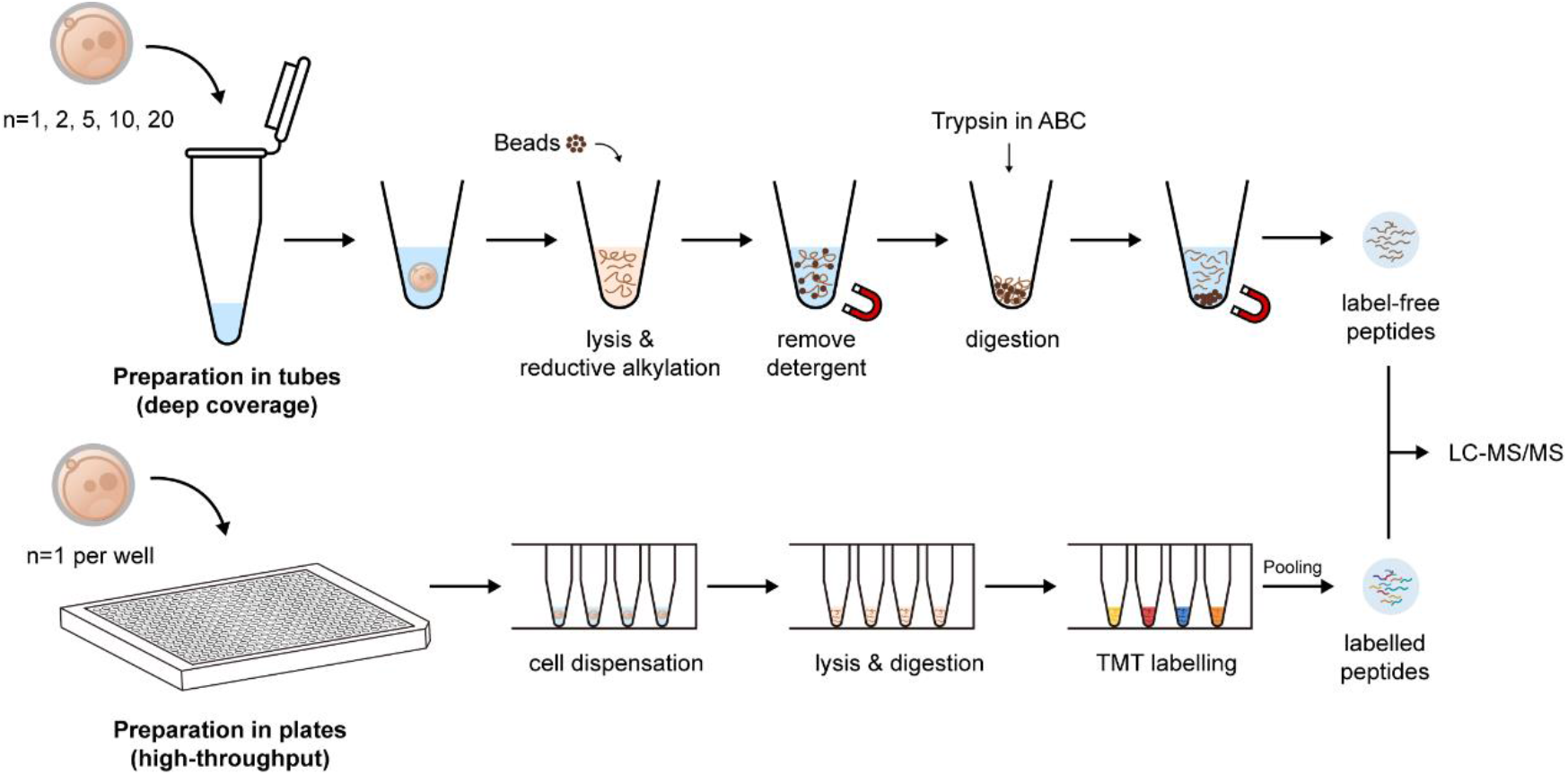
Comprehensive scheme for mouse oocyte and embryo protein profiling by ultrasensitive proteomics. The top is the deep coverage strategy which prepared in tubes without labelling. The bottom is the high-throughput strategy which prepared in 384-well plates with TMT labelling.

## Results

### Single-cell and low-input proteomics deepens zygote proteome coverage while significantly reducing starting material

Firstly, we established a full-blown proteomic workflow including one-pot sample preparation in commercial tubes and LC-MS/MS analysis based on diaPASEF mode in timsTOF (Figure 1). Then we performed this DIA-based proteomic analysis for 1, 2, 5, 10, and 20 mouse zygotes, respectively. A total of 4,700 protein groups were identified, of which an average of 2,665, 3,291, 3,893, 4,223, and 4,585 protein groups were detected in two replicates of 1, 2, 5, 10, and 20 mouse zygotes, respectively (Figure 2A). Remarkably, an average of 2,665 protein groups can be identified even in a single mouse zygote. We found the correlation of starting number of zygotes and identified protein groups number can be well fitted as a logarithm curve (R^2^=0.98), which performed better than linear fitting (R^2^=0.72) (Figure 2A). This indicated that though the number of identified protein groups increased with increasing starting materials, the benefit became progressively smaller. We also tested 30 mouse zygotes as the starting material, although their peptide level exceeded the recommended optimal detection range (≤ 200 ng) of timsTOF (Manufacturer’s recommendation). As expected, though the number of identified protein groups increased slightly, the gains were very limited (data not show). Of note, compared with searching by directDIA, protein identification can indeed benefit from a project-specific spectral library, especially as the amount of starting material increases (Figure 2B). The proportion of co-identified proteins in the two replicates of each group to the total identified proteins and the Pearson’s correlation coefficients between ten runs of zygotes revealed the excellent reproducibility and stability of proteomic data (Figure 2C-D). Of note, when the starting material increased from 1 zygote to 20 zygotes, the proportion of reproducible proteins increased from 77.8% to 96.3%. Although the proteomic data remained well reproducible even for one zygote, this also suggested that the tinier the experimental system was, the greater interferences such as instrumental or manually operated factors would disturb the results.

**Figure 2.**
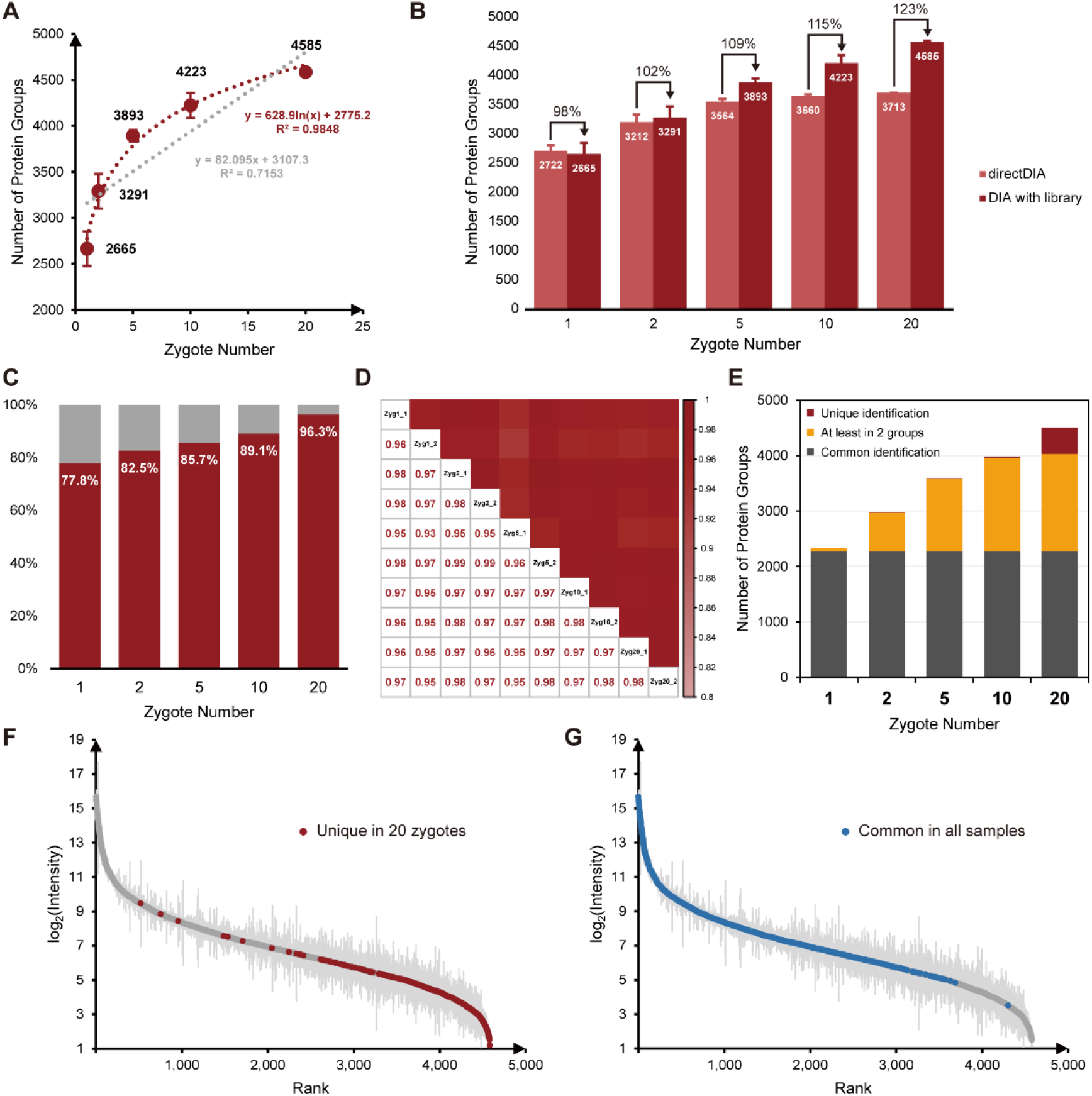
DIA-based low-input proteomics improves the zygote proteome depth while significantly reducing the starting material. (A) The number of protein groups identified from mouse zygotes of various number. Each group had two independent replicates (10 runs) and data in the plot are presented as mean ± SD. Curve Fitting was performed with R (Version 4.2.2). (B) Comparison of the number of proteins identified in each group between directDIA and spectral library-based DIA database searching. Data in the plot are presented as mean ± SD. Percentages indicate the benefit of spectral library-based protein identification. (C) The proportion of co-identified proteins in the two replicates of each group to the total identified proteins. (D) Correlation of protein expression between 10 runs of mouse zygotes (10 runs in two replicates, with five groups of different numbers of zygotes). Pearson’s correlation coefficients were presented numerically and color-coded. (E) Comparison of the number of commonly identified proteins and uniquely identified proteins between different numbers of zygotes. (F) The intensity distribution of 471 unique proteins identified in 20 mouse zygotes. (G) The intensity distribution of 2,273 proteins co-identified in five groups of different numbers of mouse zygotes.

We next compared the reproducible protein groups in different starting zygote number, finding that the 20 mouse zygotes was able to identify 471 unique proteins, mainly low-abundance proteins (Figure 2E-F). 2,273 proteins were commonly identified in all samples, covering about top 80% protein intensities (Figure 2G). GO term enrichment analysis showed that both the subcellular distribution, biological processes and molecular functions of proteins identified in different numbers of zygotes were very similar, illustrating that our characterization of the zygote proteome was unbiased (Figure 3).

**Figure 3.**
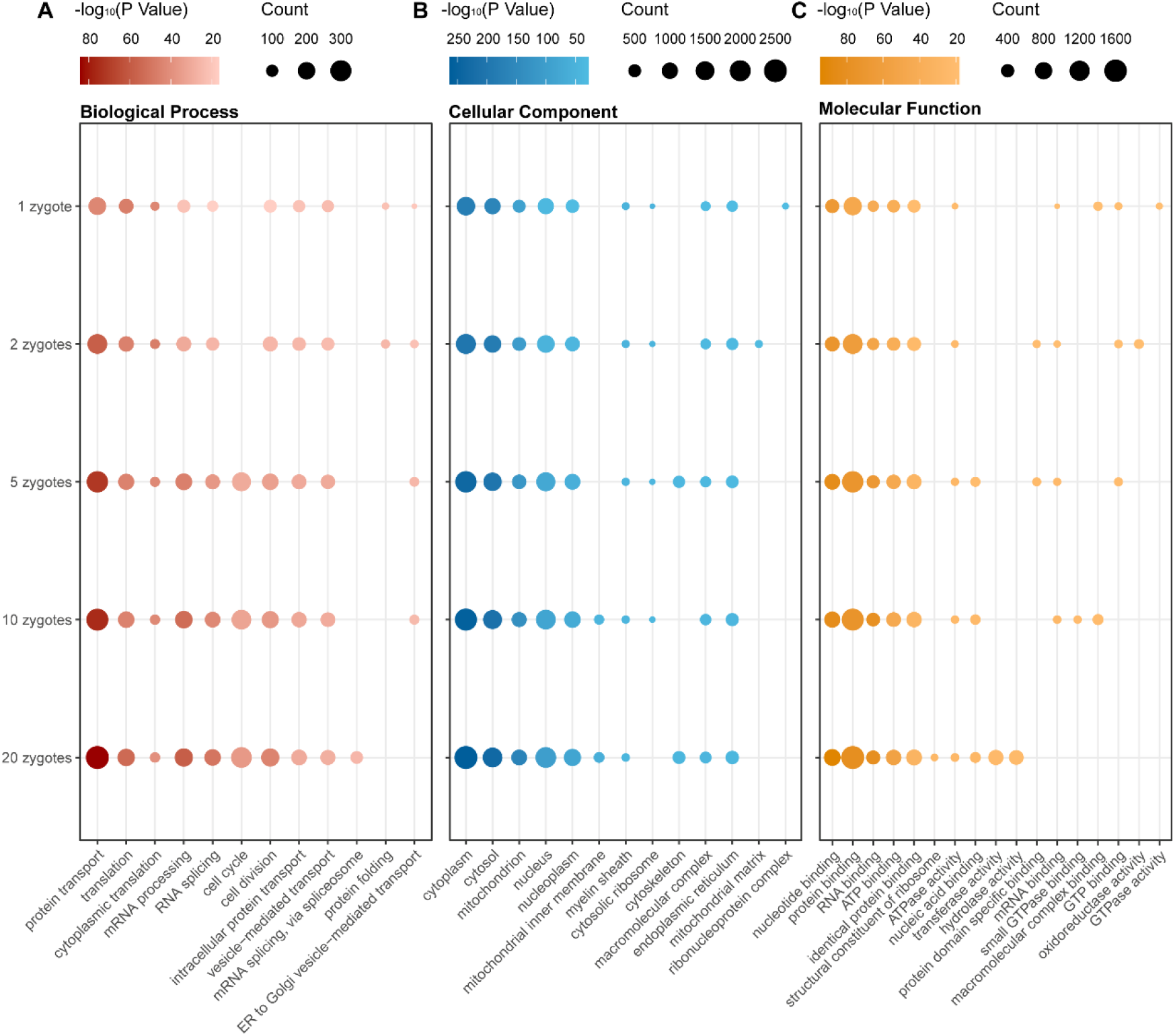
GO terms enrichment of DIA-based low-input proteomics reveals the unbiasedness of proteome from different starting materials. Dot plots of top 10 enriched GO terms including (A) biological process (BP), (B) cellular component (CC) and (C) molecular function (MF) in the identified proteins from different numbers of mouse zygotes in each group. The size of circle represents the count of protein and the color represents the -log_10_ (P value).

We also tested the same proteomic preparation workflow with DDA mode LC-MS/MS analysis, achieved an average of 1,972 and 3,437 identified protein groups from 1 and 20 mouse zygotes respectively (Figure 4A). Although the protein identification performance and data correlation were not as satisfactory as DIA mode (Figure 4B-C), these results still provided outstanding proteomic depth and can assist building project-specific spectral libraries.

**Figure 4.**
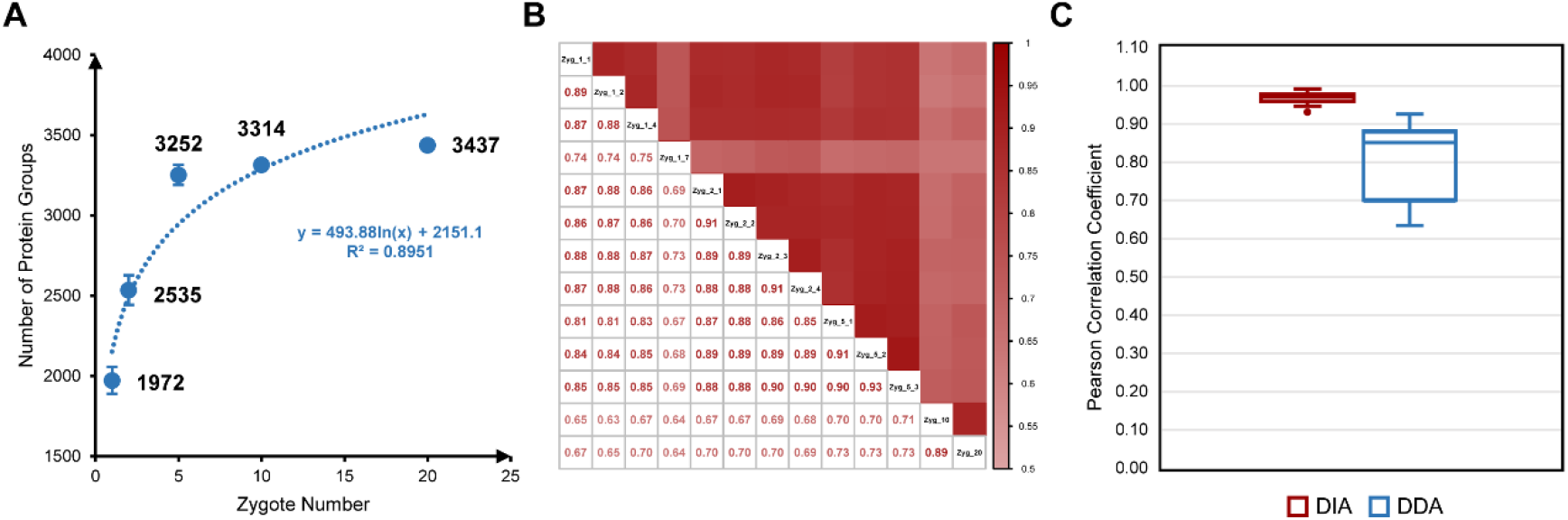
Low input proteomics based on DDA mode also has good performance in characterizing the mouse zygote proteome. (A) The number of protein groups identified from mouse zygotes of various number based on DDA mode. Data in the plot are presented as mean ± SD. Curve Fitting was performed with R (Version 4.2.2). (B) Correlation of protein expression between each run of mouse zygotes based on DDA mode. Pearson’s correlation coefficients were presented numerically and color-coded. (C) Comparison of the Pearson’s correlation coefficients between each run of the DDA and DIA mode. Boxplots show the median (middle bar), 25th and 75th percentiles (box), and 1.5× interquartile range (whiskers).

In summary, even using only one zygote starting material, the protein profile is reliable and deep enough for downstream analysis. However, if the study requires more focus on the depth of proteomic data and reproducibility of quantification, it is recommended that more starting material be used. In short, this requires balancing data integrity and quantitative reproducibility with starting material consumption.

### High-throughput proteomic profiling of mouse zygotes by multiplexed single-cell proteomics

Isobaric label-based quantification has been one of the most popular proteomic quantification strategies and been widely used for approving analysis throughput and quantitative accuracy. Tandem mass tags (TMT) are one of the most widely used isobaric labels in proteomics and have been expanded to several serials [31–33]. Slavov’s group pioneered the application of TMT to single-cell proteomics, introduced the idea of “carrier channel” which consisted by about two hundred cells to share most of the loss from single-cell channels and enhanced signal for MS analysis [34]. Several single-cell proteomic tools have been developed based on this label strategies, including the Ultra-sensitive and Easy-to-use multiplexed Single-Cell Proteomic workflow (UESCP) build in our lab [27]. As its brilliant performance in mammalian somatic cells, we tried to apply the similar strategy to single mouse zygote proteomics here. We designed a set of 6-channel experiments based on TMTsixplex (Figure 5A). Each TMT set included four channels containing one zygote in each (single-cell channel), one channel without any cell or peptide (blank channel), and one carrier channel which consisted by about 50 or 100 mouse oocytes (50× carrier or 100× carrier channel). Both 50× carrier experiment (EXP) and 100× carrier EXP were repeated twice, and an average of 2,182 or 2,371 protein groups were identified from 50x or 100x carrier EXP group (8 single mouse zygotes of each group) respectively (Figure 5B). Boxplot of protein intensities in different channels showed good data parallelism between single-cell channels, and the low intensity of blank channel confirmed the influence from carrier channel had been minimized (Figure 5C). The stabilities of defined quantitative ratios of single-cell channels illustrated that whether in 50× carrier EXP or 100× carrier EXP, the quantification of single-cell channels was reliable (Figure 5D). Though the number of proteins identified from one mouse zygote was not as many as DIA-based label-free method mentioned above, this label-based data showed higher correlation between different single zygotes (Figure 5E), which is a well-known benefit of label-based proteomics [35]. 2,021 protein groups overlapped between one zygote data from DIA-based EXP, TMT6plex 50× carrier EXP and TMT6plex 100× carrier EXP (Figure 5F), which validated the reliability of data. More importantly, multiplexed labelling increased quadruple throughput of MS analysis. Meanwhile, transferring the preparation processes from tubes to 384-well plates can excitedly enhance efficiency by more than 300-fold. To get 300 single zygotes proteomic data, the DIA-based method will take about 25 days with high labor intensity, but only about 8 days using TMT6plex label-based method. This throughput makes large-scale proteomics apply in fields of oocyte maturation and pre-implantation embryo development possible.

**Figure 5.**
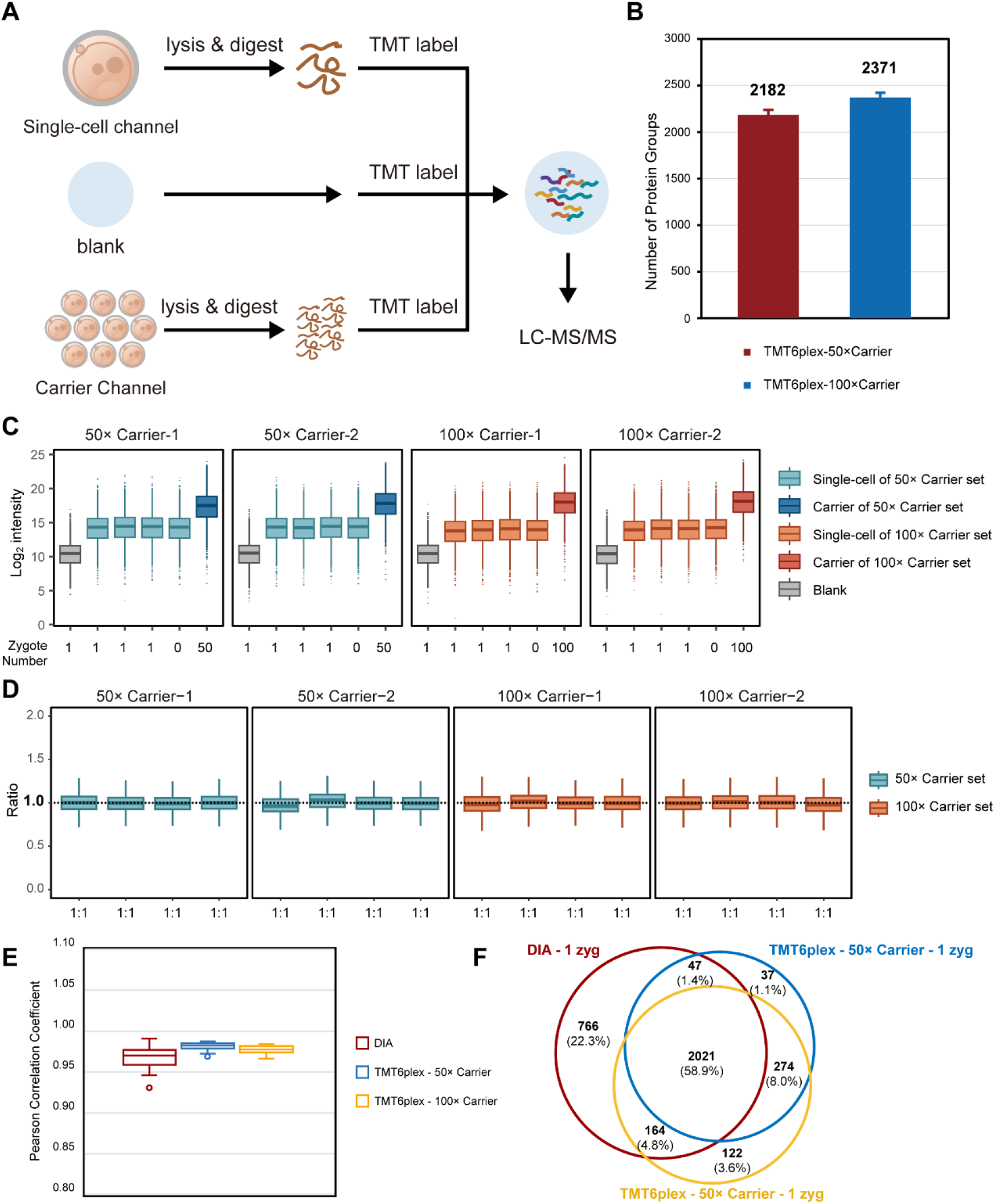
High-throughput proteomic profiling of mouse zygotes by single-cell proteomics. (A) Workflow for characterizing the mouse zygote proteome by high-throughput single-cell proteomics with the carrier channel. Each TMT6plex experiment including one blank channel (0 zygote), one carrier channel (50 or 100 oocytes), and four single-cell channels (1 zygote, respectively). (B) Comparison of the number of protein groups identified by multiplexed single-cell proteomics with TMT6plex-50× carrier EXP or 100× carrier EXP. Data in the plot are presented as mean ± SD. (C) The intensity distribution of all proteins in each TMT channel. Each different carrier TMT6plex experiment had two replicates. Boxplots show the median (middle bar), 25th and 75th percentiles (box), and 1.5× interquartile range (whiskers). (D) Boxplots of measuring ratios of protein groups based on TMT intensities between different single-cell channels in each TMT6plex experiment. The denominator is the quantitative mean of 16 sing-cell channels in four TMT experiments. (E) Comparison of the Pearson’s correlation coefficients between each run (or single-cell channel) of DIA-based proteomics, TMT6plex 50× carrier EXP and TMT6plex 100× carrier EXP. (F) Venn diagram shows the overlap between DIA-based proteomics, TMT6plex 50× carrier EXP and TMT6plex 100× carrier EXP proteomics identified proteins based on one mouse zygote.

## Discussion

Although proteomics has been a mature technique and been widely used in biological researches, few studies focused on mouse oocyte meiotic maturation and pre-implantation embryo development have been published in the last decade due to the rarity of research materials [13,21,22,24,25,36]. The development of single-cell or low-input multi-omics technologies have given new hope to this field, and have promoted the understanding of transcriptional and translational regulations during mouse preimplantation development [4,6,37]. The single-cell or low-input proteomics, however, has been still in its fledgeless stage because of the complex constituents, low abundance, wide dynamic range, and lack of amplified ability of proteins. Here we established a full set of ultrasensitive proteomic technical options for mouse embryo samples from single-cell to low-input. Using deep coverage route, an average of 2,665 protein groups can be identified in single mouse zygote, and 4,585 protein groups can be identified from 20 mouse zygotes, which has reached the depth for biological discovery. We also established the high-throughput workflow for large-scale proteomic researches. Based on TMTsixplex labelling, accomplishing the preparation and MS analysis of 300 single mouse zygotes just need 8 days, and an average of 2,182 or 2,371 protein groups can be identified from single zygote with 50× or 100× carrier.

In recent years, DIA mode has become increasingly popular because of its high data reproducibility, accuracy, and integrity [38,39]. With technological advances, TIMS has been used to separate peptide precursors based on the ion mobility (IM) dimension, and in particular the advent of PASEF has improved ion utilization, increased sensitivity, and reduced the complexity of the spectrum [40–42]. Subsequently, diaPASEF, which combines the advantages of DIA mode acquisition and the high efficiency of PASEF, was developed and applied, suitable for proteomic characterization of very limited samples [26,43–46]. Wang et al. demonstrated that diaPASEF can dramatically double the protein identification numbers compared with ddaPASEF in single mammal somatic cells [28]. Our data also showed 120% ∼ 135% improvement of protein identification using diaPASEF compared with ddaPASEF, and the higher correlation of diaPASEF proved the data reliability. A spectral library is usually important for DIA MS data analysis. Compared with library-free analysis strategy (directDIA), an appropriate spectral library can remarkably increase the quantified precursors and proteins from the same MS data [47,48]. In our work, we found that compared with directDIA, protein identification can indeed benefit from a project-specific spectral library, especially as the amount of starting material increases. These results consolidated the necessity of the appropriate spectral library for diaPASEF analysis.

Single-cell proteomics has reached a rapid developmental stage recently. Although the advances in instruments and new analysis strategies have dramatically advanced the single-cell proteomic depth, accessibility became another major inhibition for actual applications. In order to minimize the loss of trace protein during preparation, most existing single-cell or low-input proteomic routes require specialized equipment, which limits their widespread application. One of the highlights of our workflows is that they are all easy-to-use and easy to automate. Without any customized equipment, these ultrasensitive proteomic schemes can be promoted conveniently to other labs. More significantly, we have also tried to apply these workflows on human oocytes and embryos analysis, and achieved exiting results (data not show). Thus, we believe this comprehensive and easy-to-use solution will benefit more researchers in the fields about oocyte meiotic maturation or pre-implantation embryo development.

In conclusion, we gave a comprehensive solution of ultrasensitive proteome technology for single-cell and low-input mouse embryo samples. Both deep coverage and high-throughput technology routes in our ultrasensitive proteome platform could greatly improve proteome research in the field.

## Materials and methods

### Mouse

All animal experiments involving mice were performed in accordance with the guidelines of the Animal Care and Use Committee of the Institute of Neuroscience, Center for Excellence in Brain Science and Intelligence Technology, Chinese Academy of Sciences, Shanghai, China. Mice were maintained in an Assessment and Accreditation of Laboratory Animal Care credited specific pathogen-free facility under a 12 h light, 12 h dark cycle. Ambient temperature is 20 °C, relative humidity is 50%.

### Zygote collection

For zygote collection, superovulated B6D2F1 (C57BL/6J x DBA/2N) strain female mice were mated with adult B6D2F1 males [49]. For superovulation, 8-week-old female mice were injected intraperitoneally with 10 IU of pregnant mare serum gonadotropin (PMSG, San-Sheng Pharmaceutical Co. Ltd), followed by the injection of 10 IU human chorionic gonadotropin (hCG, San-Sheng Pharmaceutical Co. Ltd) 20 h later. Zygotes with two PN were collected from female oviducts and transferred from HTF medium to KSOM (Sigma-Aldric), then cultured at 37 °C under 5% CO_2_. One-cell stage embryos were collected after hCG injection for 30-32 h.

### Sample preparation in tube for DIA and DDA analysis

Clean zygotes were collected into tubes containing lysis buffer and lysed by heating [50]. TCEP and CAA were added to reduce and alkylate proteins. Then pre-washed magnetic beads (Fisher Scientific, Germany) were added to protein solution and Ethanol (EtOH) were also added to promote proteins binding. The protein-beads mixtures were incubated and then the supernatant was removed on the magnetic rack as waste. The protein-binding beads were washed to remove the detergents and other contaminants. MS-grade trypsin (Thermo Scientific™) were added to resuspend protein-binding beads. After enzymolysis, the supernatant containing digested peptides was transferred to clean tubes and acidized. Samples were then desalted in homemade C18 StageTips [51]. Clean peptides were lyophilized and resuspended in 0.1% (v/v) formate/ddH_2_O for LC-MS/MS analysis.

### Sample preparation in 384-well plate for TMT-label analysis

Single zygotes were dispensed into 384-well plates containing lysis buffer. The plates were frozen and heated to lysis cells and release proteins by freeze-thaw. MS-grade trypsin were added and the 384-well plates were incubated at 37 °C to digest proteins. Peptides were then labelled with TMTsixplex™ reagents (Thermo Fisher Scientific), and then followed by quenching. Differentially labelled samples from one TMT set were combined and desalted in homemade C18 StageTips. Clean peptides were lyophilized and resuspended in 0.1% (v/v) formate/ddH_2_O for LC-MS/MS analysis.

For TMT6plex set, the single zygotes were labelled with different channels. The carrier sample was prepared in bulk through the filter assisted sample preparation (FASP) procedure [52] which used 1,000 oocytes as starting material. Peptides equivalent of 50 or 100 oocytes were prepared as same steps as single-cell samples synchronously, labelled with the other TMT channel. Blank samples were also prepared as same steps as single-cell samples synchronously without any cell or peptide adding as starting material, and labelled with another TMT channel.

### LC-MS/MS analysis

All samples were separated via a high performance applied chromatographic system nanoElute® (Bruker). Peptides were loaded on to an in-house packed column (75 μm × 250 mm; 1.9 μm ReproSil-Pur C18 beads, Dr. Maisch GmbH, Ammerbuch) which was heated to 60 °C, and separated with a 60-min gradient of 2% to 80% mobile phase B at a flow rate of 300 nL/min. The mobile phases A and B were 0.1% (v/v) formate/ddH_2_O and 0.1% (v/v) formate/acetonitrile, respectively.

Mass spectrometry analysis was performed on a trapped ion mobility spectrometry coupled to time-of-flight mass spectrometer (timsTOF MS, Bruker). Parallel accumulation-serial fragmentation (PASEF) was applied in all sample analysis [53]. MS and MS/MS spectra were recorded from m/z 100 to 1,700. For TMT-label samples and label-free samples which acquired by data-dependent acquisition (DDA) mode, one topN acquisition cycle took 100 ms and contained one MS1 survey TIMS-MS and 8 PASEF MS/MS scans with two TIMS stepping. For label-free samples which acquired by data-independent acquisition (DIA), we set an acquisition scheme with 64 precursor isolation windows.

### Database searching

Mass spectrometry raw files of TMT-label experiments and DDA label-free experiments were processed with PEAKS software (Version Online X) and searched against the UniProt/SwissProt mouse database (UP000000589, containing 17,090 entries, downloaded on Nov 24, 2021) using the default factory settings. In brief, Trypsin/P was set as the digestion enzyme with maximum two mis-cleavage sites allowed. False-discovery rates were controlled at 1% both on peptide and protein group levels. Only peptides with a length from 7 to 52 amino acids were considered for search. The oxidation of methionine and acetylation of protein N-term were set as variable modifications, and carbamidomethyl of cysteine was set as fixed modification.

DIA label-free data was processed with Spectronaut® (v16; Biognosys). For directDIA search, raw files were searched against the above UniProt/SwissProt mouse database as same as DDA analysis with default params. For library-based DIA search, raw files were searched against a self-established hybrid spectral library with default params.

### Data analysis

Statistical analyses and visualizations were performed using R (Version 4.2.2) unless otherwise stated. Data are presented as mean value ± standard deviation (SD). Protein gene ontology (GO) term enrichment analysis was achieved by The Database for Annotation, Visualization and Integrated Discovery (DAVID)(v2022q3) [54,55].

## CRediT author statement

Conceptualization, Z.Y.L., Z.L., C.L., and H.W.; Methodology & Investigation, L.G., X.M.L., W.C.Z., Y.S., Z.Y.L., W.Q.Q., Z.H.P., and L.J.Q.; Writing – Original Draft, C.L, L.G., X.M.L., Q.Q.W., and Z.Y.L.; Writing – Review & Editing, W.C.Z., Q.Q.W., H.P.Z., and L.J.Q.; Supervision, Z.Y.L., Z.L., C.L., and H.W.; All authors contributed to the interpretation of the data and read and approved the final manuscript.

## Acknowledgments

We thank all members of our laboratories for helpful discussions. This work was supported by the National Natural Science Foundation of China (NSFC) (grants 82030099, 30700397), the Shanghai Municipal Science and Technology Commission “Science and Technology Innovation Action Plan” technical standard project (21DZ2201700), the Shanghai Municipal Science and Technology Major Project (2018SHZDZX05), the Strategic Priority Research Program of the Chinese Academy of Sciences (XDB32060105), the Basic Frontier Scientific Research Program of CAS (ZDBS-LY-SM019), and the innovative research team of high-level local universities in Shanghai.

